# Characterization of GAPDH expression related to biotic stress responses in Physcomitrella

**DOI:** 10.1101/2023.06.07.544018

**Authors:** Alessandra A. Miniera, Sebastian N.W. Hoernstein, Nico van Gessel, Jens O. Peeters, Ralf Reski

**Affiliations:** Plant Biotechnology, Faculty of Biology, University of Freiburg, Schänzlestr. 1, 79104 Freiburg, Germany; CIBSS –Centre for Integrative Biological Signalling Studies, Schänzlestr. 18, 79104 Freiburg, Germany

**Keywords:** hormonal regulation, immunity, methyl jasmonate, moss, pathogens, salicylic acid

## Abstract

Glyceraldehyde 3-phosphate dehydrogenase (GAPDH) is a housekeeping protein that plays an important role in the energy metabolism and is found in all kingdoms of life. While animals possess one GAPDH isoform, plants possess several GAPDHs. GAPA and GAPB are chloroplastic phosphorylating GAPDHs involved in the Calvin-Benson cycle. GAPC in the cytosol and GAPCp in plastids are glycolytic phosphorylating GAPDHs involved in the glycolysis pathway. In animals as well as in plants, GAPDH isoforms have various non-metabolic roles, e.g. in autophagy, apoptosis, and were identified as redox sensors. In plants, in addition to the variety of moonlight functions in abiotic stress, GAPDHs are also involved in biotic stress responses. Here, we identified 17 GAPDH isoforms from the moss Physcomitrella, belonging to the *GAPA*, *GAPC,* and *GAPCp*. We searched for protein and transcript evidences in publicly available proteome and transcriptome data, laying the focus on datasets of treatments with the phytohormones salicylic acid (SA) and methyl jasmonate (MeJA). We investigated the putative role of GAPDHs in plant immune response by identifying SA- and MeJA-inducible *GAPA*s and *GAPC* genes. The *GAPA*s Pp3c1_13170 and Pp3c11_15790 and the *GAPC* Pp3c18_15450 appear to be unresponsive to SA treatment. In contrast, we identified four MeJA-responsive genes. MeJA-treatment resulted in a 10- and 3-fold downregulation of the *GAPA*s Pp3c1_13170 and Pp3c11_15790, whereas expression of the two *GAPC* genes Pp3c18_15450 and Pp3c21_9380 showed an up to 7- and 33-fold upregulation after 4 hours of MeJA treatment, respectively. Simultaneously, a four-hour MeJA-treatment induced the remodeling of the Physcomitrella secretome, resulting in the production of antimicrobial compounds, which in turn led to a bacteriostatic growth inhibition of 26% of *E. coli.* These findings draw attention to the potential differential regulation of GAPDH genes in plant immune response as well as a potential role for GAPC in the defense response against necrotrophic pathogens.

## Introduction

Plants are subjected to a variety of pathogens, such as bacteria, viruses, fungi as well as insects and nematodes. Pathogens can be categorized into three groups, namely biotrophs, necrotrophs or hemi-biotrophs. Biotrophic pathogens feed on the living plant tissue, while necrotrophic pathogens first destroy the plant tissue before feeding on the nutrients. Pathogens which perform both the biotrophic and necrotrophic life style are referred to as hemi-biotrophic pathogens (Pieterse *et al*., 2012). Unlike animals, plants solely rely on their innate immune system (Jones & Dangl, 2006). Two pathogen response mechanisms are induced upon successful entrance of pathogens, namely the microbial-triggered immunity (MTI) and the effector-triggered immunity (ETI) (Muthamilarasan & Prasad, 2013). Microbe-associated molecular patterns (MAMPs) or damage-associated molecular patterns (DAMPs) are recognized by pattern recognition receptors (PRR) which activate the MTI response. Successful pathogens employ effectors which counteract the MTI response, resulting in an effector-triggered immunity (ETI). The ETI response usually leads to programmed cell death (PCD) referred to as hypersensitive response, which is observable at the infection site (Jones & Dangl, 2006).

Both pathogen-response mechanisms lead to significant cellular changes such as Ca^2+^ flux, production of nitrogen intermediates (e.g. *via* S-nitrosylation of cysteine residues) and reactive oxygen species (ROS), activation of the MAPK cascade, expression of defense genes, the biosynthesis of the phytohormones jasmonic acid (JA) and salicylic acid (SA) (Muthamilarasan & Prasad, 2013) and production of antimicrobial compounds (Wang *et al*., 2022).

Salicylic acid, JA and ethylene (ET) are key regulators of the innate immune system and the SA-JA interaction is mostly of an antagonistic nature, but can also be synergistic or neutral (Pieterse *et al*., 2012). An example of their antagonistic interaction is their effect on the cellular redox buffer glutathione. SA increases and JA decreases the glutathione pool (Pieterse *et al*., 2012). While SA is a positive regulator of immune responses against biotrophic pathogens, JA is a positive regulator of immune responses against necrotrophic pathogens and is associated with wounding and defense responses against insects. The phytohormones abscisic acid (ABA), cytokinin (CK), auxin (AUX), gibberellic acid (GA) and brassinosteroids (BR) are associated with plant development and abiotic stress responses but are also known to fine-tune either the SA- or JA/ET immune response pathways (Shigenaga & Argueso, 2016). In Physcomitrella, the presence of AUX, CK, ABA, SA, 12-oxo-phytodienoic acid (OPDA, JA precursor) and strigolactones (SL) have been described (Guillory & Bonhomme, 2021; Bennett *et al*., 2014; Lüth *et al*., 2023; Lindner *et al*., 2014; Stumpe *et al*., 2010; Arif *et al*., 2019; Decker *et al*., 2017). Despite Physcomitrella not being able to produce JA, it can respond to MeJA treatment and SA treatment with transcript accumulation of the defense gene phenylalanine ammonia-lyase (PAL) (Ponce De León *et al*., 2012). SA and MeJA treatments caused the alteration of intracellular and secretome peptide pools by inducing the degradation of protein precursors (Filippova *et al*., 2019; Fesenko *et al*., 2019). Additionally, MeJA treatment of Physcomitrella resulted in the production of antimicrobial peptides from functional precursors (Fesenko *et al*., 2019).

Glyceraldehyde 3-phosphate dehydrogenase (GAPDH) is known as housekeeping protein for its role in the glycolysis pathway and it is conserved across all kingdoms of life. Animals possess one GAPDH isoform and recent studies described various moonlighting non-metabolic functions of GAPDH such as RNA-and telomere-binding, DNA transcription, DNA repair, cell-cycle regulation, membrane fusion and trafficking, protein binding, autophagy and apoptosis (Colell *et al*., 2007; Colell *et al.,* 2009; Tarze *et al*., 2007; Tristan *et al*., 2011; Zhang *et al*., 2015). Several post-translational modifications (PTMs; e.g.: phosphorylation, acetylation, carbonylation, S-nitrosylation, S-thiolation) at conserved amino acid residues have an effect on cellular localization and functions of GAPDH (Zhang *et al*., 2015; Butterfield *et al*., 2010; Tristan *et al*., 2011). Especially the conserved catalytic cysteine was shown to carry various PTMs. S-nitrosylation of the catalytic cysteine activated the GAPDH binding to the E3-ligase Siah1, which facilitated the degradation of nuclear proteins and induction of apoptosis in mouse cell lines (Hara *et al*., 2005). Oxidative stress in human cells lines showed a protein-protein interaction of GAPDH with the MAP kinase JNK (Itakura *et al*., 2023). Plants possess several GAPDHs which are located at different cell compartments and can be separated into two major groups: phosphorylating and non-phosphorylating GAPDHs. GAPA/B, GAPC and GAPCp are (de)phosphorylating GAPDHs which act either in the Calvin-Benson cycle (GAPA/B) or glycolysis (GAPC in cytosol and GAPCp in plastids). *GAPB* and *GAPCp* resulted from gene duplications of *GAPA* and *GAPC* respectively (Petersen *et al.,* 2003; Petersen *et al*., 2006; Martin & Cerff, 2017). GAPA/B catalyze the NADPH-specific dephosphorylation of 1,3-bisphosphoglyceric acid (BPGA) to glyceraldehyde-3-phosphate (G3P). The glycolytic GAPC and GAPCp catalyze the strictly NAD^+^-dependent reversible phosphorylation of G3P to BPGA (Zaffagnini *et al*., 2013). GAPN is the sole member of the non-phosphorylating group which catalyzes the oxidation of G3P directly to 3-phosphoglycerate (3PGA), consequently bypassing the intermediate reaction catalyzed by GAPC (Habenicht *et al.,* 1994). Similar to the GAPDH isoform in animals, cytoplasmic GAPC from different plants and their catalytic cysteine were reported to be target for various redox post-translational modification, such as sulfonation, glutathionylation and S-nitrosylation, resulting in the inhibition or alteration of GAPC activity (Zaffagnini *et al*., 2013). Studies in Arabidopsis, potato, rice, wheat and sheepgrass have described the role of GAPDH isoforms in abiotic stress responses such as heat, cold, drought, alkali- and salt stress (Hildebrandt *et al*., 2015; Yang *et al*., 1993; Guo *et al*., 2012; Li *et al*., 2019; Lu *et al*., 2020; Kim *et al.,* 2020). GAPC is described as an H_2_O_2_ sensor and was shown to be very sensitive to oxidative stress inhibiting its activity (Yang & Zhai, 2017). All phosphorylating GAPDHs in Arabidopsis are redox sensitive (Henry *et al*., 2015). GAPDH also plays a role in H_2_O_2_-mediated apoptosis, as the overexpression of GAPA1 in Arabidopsis protoplasts resulted in the inhibition of ROS production and BAX-induced apoptosis (Baek *et al*., 2008). Plant GAPDH isoforms also play a role in biotic stress responses. In potato leaves transcript levels of GAPC were increased upon infection with the fungus *Phytophthora infestans* and elicitor treatment (Laxalt *et al*., 1996). In Arabidopsis immune elicitation with flg22 showed an increase in intracellular ROS in *GAPC* KO mutants and all *GAPDH* KO lines showed an enhanced HR and accelerated apoptosis in response to ETI. Additionally, the treatment with flg22 showed the contrasting regulation of the *GAPA*s, *GAPB* and *GAPC1* during MTI (Henry *et al*., 2015). A GAPDH-derived peptide (YFGAP) was isolated from the yellowfin tuna *Thunnus albacares* showing a bacteriostatic antimicrobial activity against gram-negative and gram-positive bacteria (Seo *et al*., 2012). In Physcomitrella, MeJA treatment resulted in a GAPDH-derived peptide (Pp3c2_24160) which showed antimicrobial activity by inhibiting bacterial growth (Fesenko *et al*., 2019).

Here, we tested the involvement of Physcomitrella GAPDHs from different subfamilies in plant immune response by identifying SA- and MeJA-inducible *GAPDH* genes. Transcript analyses revealed that while *GAPA*s seem to be unresponsive to SA- and downregulated by MeJA-treatment, two *GAPC* genes were identified as MeJA-inducible genes. These results highlight the possible distinct regulation of *GAPDH* genes in plant immune response and a putative involvement in GAPC in defense responses against necrotrophic pathogens.

## Results

### Phylogenetic analysis of phosphorylating GAPDH genes

In our work we focused on the phosphorylating GAPDHs. Using the GAPA from Arabidopsis (At1G12900) as query we performed proteome-wide searches against the latest protein models of six selected plant species: *Physcomitrium patens* (V3.3; Lang *et al.,* 2018, *Sphagnum angustifolium* (V1.1; Healey *et al*., 2023), *Marchantia polymorpha* (V3.1; Bowman *et al*., 2017), *Arabidopsis thaliana* (Araport11; Cheng *et al*., 2017), and *Oryza sativa* (V7.0; Ouyang *et al*., 2007), all available from the Phytozome13 database (Goodstein *et al*. 2012; https://phytozome-next.jgi.doe.gov/), and complemented by *Funaria hygrometrica* (Kirbis *et al*., 2020). In a recent publication (Healey *et al*., 2023) it was clarified that *Sphagnum angustifolium* had originally been misclassified and wrongly annotated as *Sphagnum fallax*. Since to this date no updated accession IDs have been available on Phytozome, we kept the original gene IDs for *S. angustifolium* in the following graphics. In total, we identified 61 protein sequences corresponding to distinct genes and whose homology was confirmed by reciprocal best hit searches. Following our phylogenetic reconstruction (Figure 1) we resolved the four clades containing 21 *GAPA*, 3 *GAPB*, 23 *GAPC*, and 14 *GAPCp* genes. Remarkably, while the majority of the selected species encode for 5-7 phosphorylating GAPDHs, our analysis revealed several additional GAPDH genes in Physcomitrella and *F. hygrometrica*, both belonging to the moss family of Funariaceae. In total we identified 17 phosphorylating GAPDHs in Physcomitrella, belonging to the *GAPA* (6), *GAPC* (8), and *GAPCp* (3) clades whereas in *F. hygrometrica* 18 were identified (8 *GAPA*, 7 *GAPC*, 3 *GAPCp*). Surprisingly, our phylogenetic analysis revealed six additional homologs in Physcomitrella and five in *F. hygrometrica*, respectively, forming a distinct subclade of GAPC and sharing predicted cytoplasmic localization (Supplementary Table 1). While almost all *GAPA/B* and *GAPC* and *GAPCp* genes are highly expressed across multiple tissues, all Funariaceae-exclusive *GAPC* genes are hardly expressed in any of the developmental stages of Physcomitrella (Supplementary Figure 1). In addition, the two Funariaceae-exclusive GAPCs Pp3c24_16410V3.1 and Funhy_Fh_22091 lack the highly conserved catalytic cysteine. Hence, we consider these isoforms as non-active regarding their glycolytic activity.

**Figure 1:**
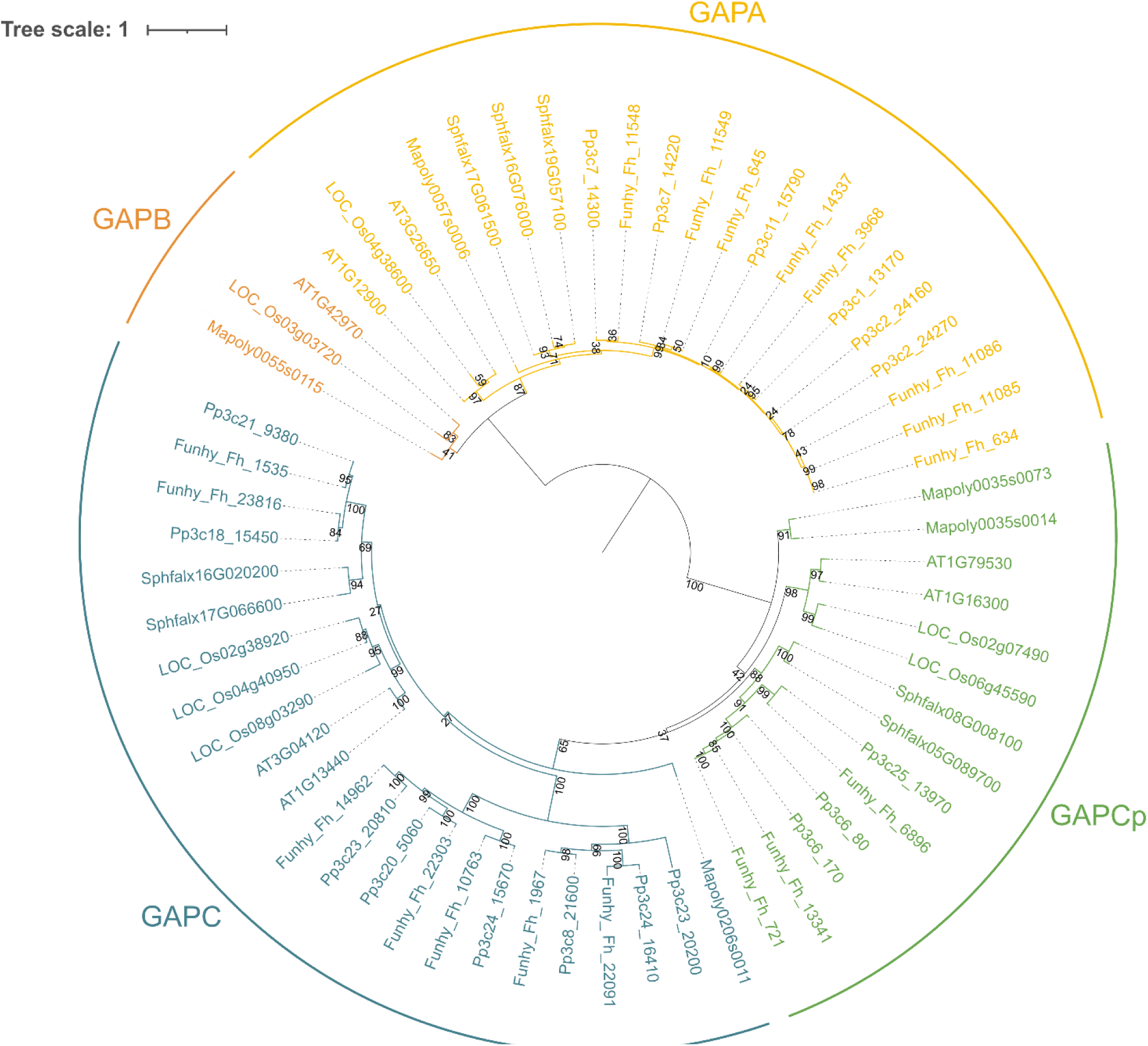
Phylogenetic tree of phosphorylating GAPDHs from selected plant species. **The analysis of** 61 phosphorylating GAPDH protein sequences from six plant species resolved four subclades containing 21 GAPAs, 3 GAPBs, 23 GAPCs, and 14 GAPCps. In total we identified 17 GAPDH-encoding genes in Physcomitrella belonging to the *GAPA*, *GAPC* and *GAPCp* subclades. Species abbreviations: Pp: *Physcomitrium patens*, Fh: *Funaria hygrometrica*, Sphfalx: *Sphagnum angustifolium*, Mapoly: *Marchantia polymorpha*, Os: *Oryza sativa*, At: *Arabidopsis thaliana*.

### Proteomic and transcriptomic evidence of GAPDHs

To investigate whether the 17 phosphorylating GAPDHs could be involved in pathogen response in Physcomitrella we inspected publicly available proteome and transcriptome data and searched for protein and transcript evidence. The focus was laid on datasets generated from treatments with the phytohormones salicylic acid (SA) or jasmonic acid (JA). The phytohormones salicylic acid (SA) and jasmonic acid (JA) are among the key regulators of plant immunity (Bari & Jones, 2009). SA treatment in Physcomitrella led to a remodeling of the extracellular peptidome such as decreasing the pool of modified peptides (Filippova *et al*., 2019). Likewise, treatment with MeJA lead to a remodeling of the intracellular proteome, the secretome and the extracellular peptidome (Fesenko *et al*., 2019). Thus, we compiled existing proteome and peptidome datasets to gather information about protein abundance or proteolytic fragments with AMP potential for the 17 GAPDHs. For this we used 10 mass spectrometry (MS) datasets available from ProteomeXchange (http://www.proteomexchange.org; Deutsch *et al*., 2023; dataset identifiers are listed in Supplementary Table 2).

These datasets comprise analyses of various phytohormone (SA, MeJA) treatments, and different tissues and samples types (gametophores, protonema, protoplasts, and secretome samples) (Supplemental Table 2). The MS datasets which comprised around 160 GB of MS raw data, were again processed (Mascot Distiller, V2.7.10) and searched against the Physcomitrella protein database (Version 3.3, Lang *et al*., 2018). Relevant post-translational modifications such as N-terminal acetylation, proline hydroxylation, tyrosine/serine/threonine sulfation (among other common PTMs) were included into the search parameters. Using a custom designed bioinformatic pipeline we searched for ID, and (partial) sequence matches with the previously identified PpGAPDH candidates in the resulting identification lists.

In total we identified 27,358 peptides from 2,539 proteins, from which 239 peptides were assigned to 10 PpGAPDHs. We found proteomic evidence for all GAPAs, two GAPCps (Pp3c6_80, Pp3c170) and two GAPCs (Pp3c21_9380, Pp3c18_15450; summarized in Table 1). Of particular interest were candidates found in datasets of phytohormone treatments, secretome and peptidome analyses. These proteomic evidences can support the idea of an involvement in pathogen response, and the secretion of GAPDH-derived peptides. Almost all GAPAs had proteomic evidences with peptides found in MeJA and/or SA treated samples, as well as secretome and peptidome samples (Table 1).

**Table 1:**
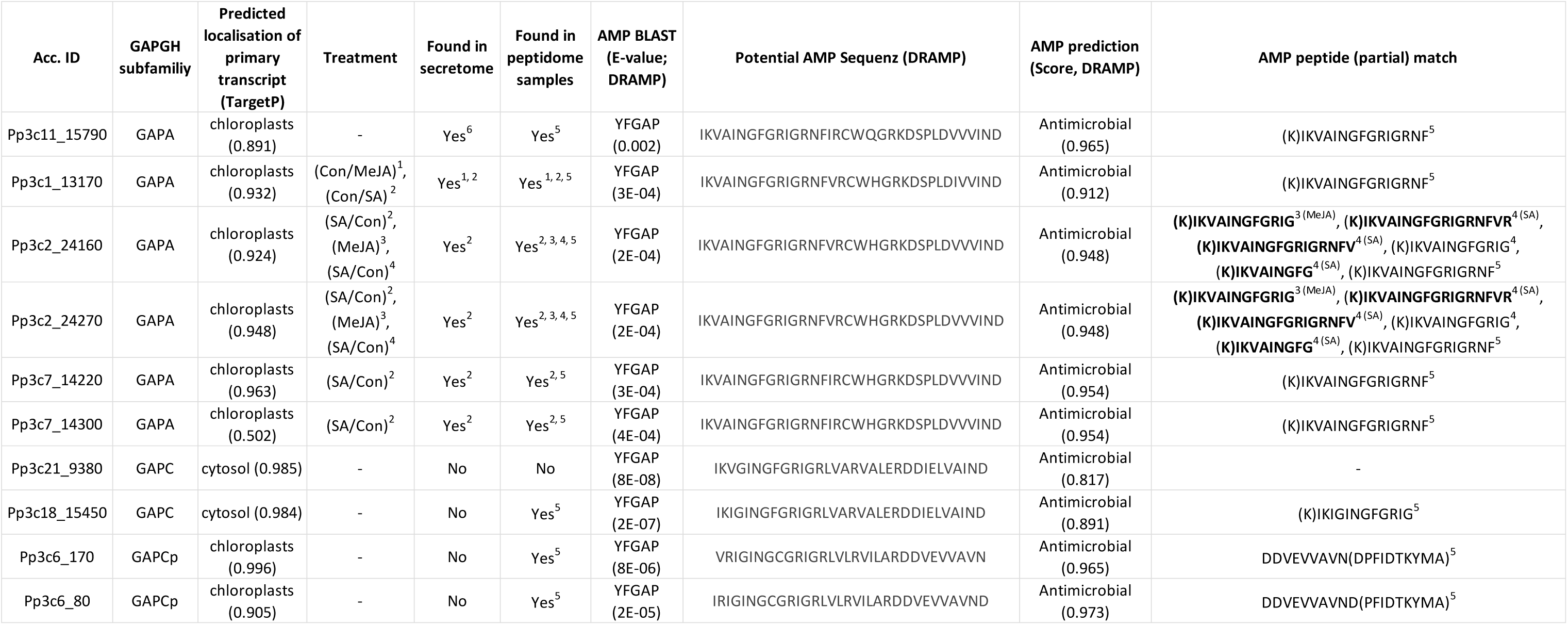
Summary of proteomic and transcriptomic evidence of identified GAPDH proteins from Physcomitrella. Across the 10 compiled MS datasets we identified 10 of the 17 PpGAPDH isoforms. Of particular interest were candidates found in datasets of phytohormone treatments, secretome and peptidome analyses. Datasets: (1) Secreted peptidome analyses of MeJA treated samples (Fesenko *et al*., 2019), (2) Secreted peptidome analyses of SA treated samples (Filippova *et al*., 2019), (3) Cellular peptidome analysis of MeJA treated protonema samples (Fesenko *et al.,* 2019), (4) Cellular peptidome analysis of SA treated protonema samples (Fesenko *et al.,* 2019), (5) Cellular peptidome analysis of protonema, protoplasts and gametophore samples (Fesenko *et al*., 2015), (6) Secreted proteome analyses of Chitosan treated samples (Lehtonen *et al*., 2013).

All GAPAs, one GAPC (Pp3c18_15450) and both GAPCps were identified in peptidome samples. Additionally, 5 of the 6 GAPAs were identified in secretome samples. The GAPC encoded by the gene Pp3c21_9380 was exclusively found in one protonema proteome dataset, however publicly available transcriptomic data shows its high potential, due to its increased expression upon OPDA (12-oxo-phytodienoic acid, MeJA precursor) treatment (Supplementary Figure 2).

Interestingly, all of the identified GAPDHs contain a sequence motif with close similarity to the antimicrobial peptide YFGAP from *Thunnus albacares* (Table 1). Moreover, an independent prediction for bioactive peptide sequences using DRAMP (Shi *et al*., 2022) confirmed the presence of a putative AMP sequence in all GAPDHs conveying activity against gram-negative bacteria (Table 1). Using those predicted AMP sequences, we reinvestigated the compiled MS data for full or partial sequence matches. We searched for proteomic evidence of the derived putative AMP sequence in our peptidome database and found several partial sequence matches for 9 of the 10 identified GAPDHs (Table 1). Interestingly, several partial sequence hits were found for the two GAPAs encoded by the genes Pp3c2_24160 and Pp3c2_24270 (Table 1). Several peptides (indicated in bold letters) were found exclusively in phytohormone-treated samples, but not in the corresponding non-treated control (Table 1). The peptides KIKVAINGFGRIGRNFVR, KIKVAINGFGRIGRNFV and KIKVAINGFG were exclusively found in SA treated proteome samples. The peptide KIKVAINGFGRIG was exclusively found in MeJA treated samples (Table 1).

In addition, we checked for transcriptomic evidence using publicly available data (PEATmoss; Fernandez-Pozo *et al*., 2020; https://peatmoss.plantcode.cup.uni-freiburg.de). The expression of the Physcomitrella *GAPDH* genes in the different developmental stages greatly differ among the GAPDH subfamilies as well as within the subfamilies (Supplementary Figure 1). All *GAPA*s are expressed in germinating spores, protonema, gametophores, leaflets and protoplasts, while they are hardly expressed in dry and imbibed spores. Most of the *GAPA* genes present a stronger expression compared to *GAPC* and *GAPCp* genes. The *GAPA*s Pp3c1_13170 and Pp3c11_15790 show the highest expression among the *GAPA* subgroup, while the *GAPA*s Pp3c2_24160 and Pp3c7_14220 show the lowest expression. Among the *GAPC*s exclusively the genes Pp3c21_9380 and Pp3c18_15450 show a high expression, with Pp3c18_15450 being expressed in all tissues, and Pp3c21_9380 is highly expressed exclusively in dry and imbibed spores. The *GAPC*s homologs which were found in the Funariaceae-exclusive subclade (Pp3c20_5060, Pp3c23_20200, Pp3c23_20810, Pp3c24_15670, Pp3c24_16410 and Pp3c8_21600) are hardly expressed in any developmental stages of Physcomitrella. Among the *GAPCp*s the gene Pp3c6_170 shows the highest expression across all tissues, while Pp3c6_80 shows a lower expression and Pp3c25_13970 is hardly expressed in any of the developmental stages.

Additionally, we referred to the publicly available expression data (PEATmoss) showing the gene regulation upon OPDA treatment (Supplementary Figure 2). Exclusively the *GAPC* Pp3c21_9380 is upregulated (3-fold) upon OPDA treatment, rendering this gene a suitable candidate for testing of its MeJA-responsivity.

### Gene expression analysis of selected GAPDHs in Physcomitrella

To further elucidate the putative role of GAPDHs from different subclades in plant immunity we analyzed the gene expression of selected candidates in response to SA- and MeJA-treatment *via* qRT-PCR.

We selected candidates from different subclades based on their proteomic and transcriptomic evidence. For all except one GAPDH (Pp3c21_9380) we found proteomic evidence either in secretome or peptidome samples (Table 1). This is especially interesting since none of the 10 candidates is predicted to be secreted to the extracellular space and hence, this could indicate a specific function in an antimicrobial response. This is further supported by the fact the proteolytic fragments with potential antimicrobial activity of several GAPDHs (e.g. Pp3c11_15970, Pp3c1_13170) were identified in peptidome samples. For Pp3c21_9380 no proteomic evidence was identified in any analyzed datasets of MeJA and SA treatments. However, according to publicly available expression data, a 3-fold increase in gene expression in response to the JA precursor OPDA was detected and hence this gene was included in the expression analysis performed in the present study (Supplementary Figure 2). For some of the 10 candidates no primers satisfying the requirements could be designed which was mostly due to off-target effects related to high sequence similarities across the isoforms. Finally, we selected the two *GAPA*s Pp3c11_15790 and Pp3c1_13170 and two *GAPC*s Pp3c21_9380 and Pp3c18_15450 for which appropriate primers for expression analysis could be obtained. To get a first impression on expression level changes we performed the phytohormone treatments of protonema suspension culture over the course of 24 hours. For this purpose, we performed the qRT-PCR analysis first in technical replicates. Because Ponce De León *et al*. (2012), Filippova *et al*. (2019) and Fesenko *et al*. (2019) reported that 400-500 µM SA and 400 µM MeJA were sufficient to elicit a response of Physcomitrella on proteome, transcriptome or phenotypical level, we performed phytohormone treatments using 400 µM SA or MeJA, respectively.

The *GAPA* Pp3c1_13170 was significantly upregulated about 3-fold compared to the untreated Mock sample after 4 hours treatment with 400 µM SA (Figure 2A). In contrast, the gene was downregulated about 2-fold after 24 hours of treatment. The *GAPA* Pp3c11_15790 and the *GAPC* Pp3c18_15450 showed no significant alteration in expression during the SA treatment (Figure 2A). In contrast, treatment with 400 µM MeJA induced significant changes in gene expression for all four GAPDHs analyzed here. The two plastidic *GAPA*s (Pp3c1_13170, Pp3c11_15790, Figure 2B) were strongly downregulated after MeJA treatment. Here, Pp3c1_13170 was downregulated by 2.6-fold after 1 hour and about 10-fold after 4 and 24 hours of MeJA treatment. The *GAPA* Pp3c11_15790 was significantly downregulated about 3-fold after 4 and 24 hours. The cytosolic *GAPC*s (Pp3c18_15450, Pp3c21_9380, Figure 2B) showed a strongly increased gene expression after all three time points of MeJA treatment. Pp3c18_15450 was upregulated about 4-fold after 1 and 24 hours with a maximum of 7.6-fold after 4 hours of treatment. The *GAPC* Pp3c21_9380 showed the strongest increase in gene expression with a 16-fold upregulation after 1 hour and a 33-fold upregulation after 4 and 24 hours of the MeJA treatment.

**Figure 2:**
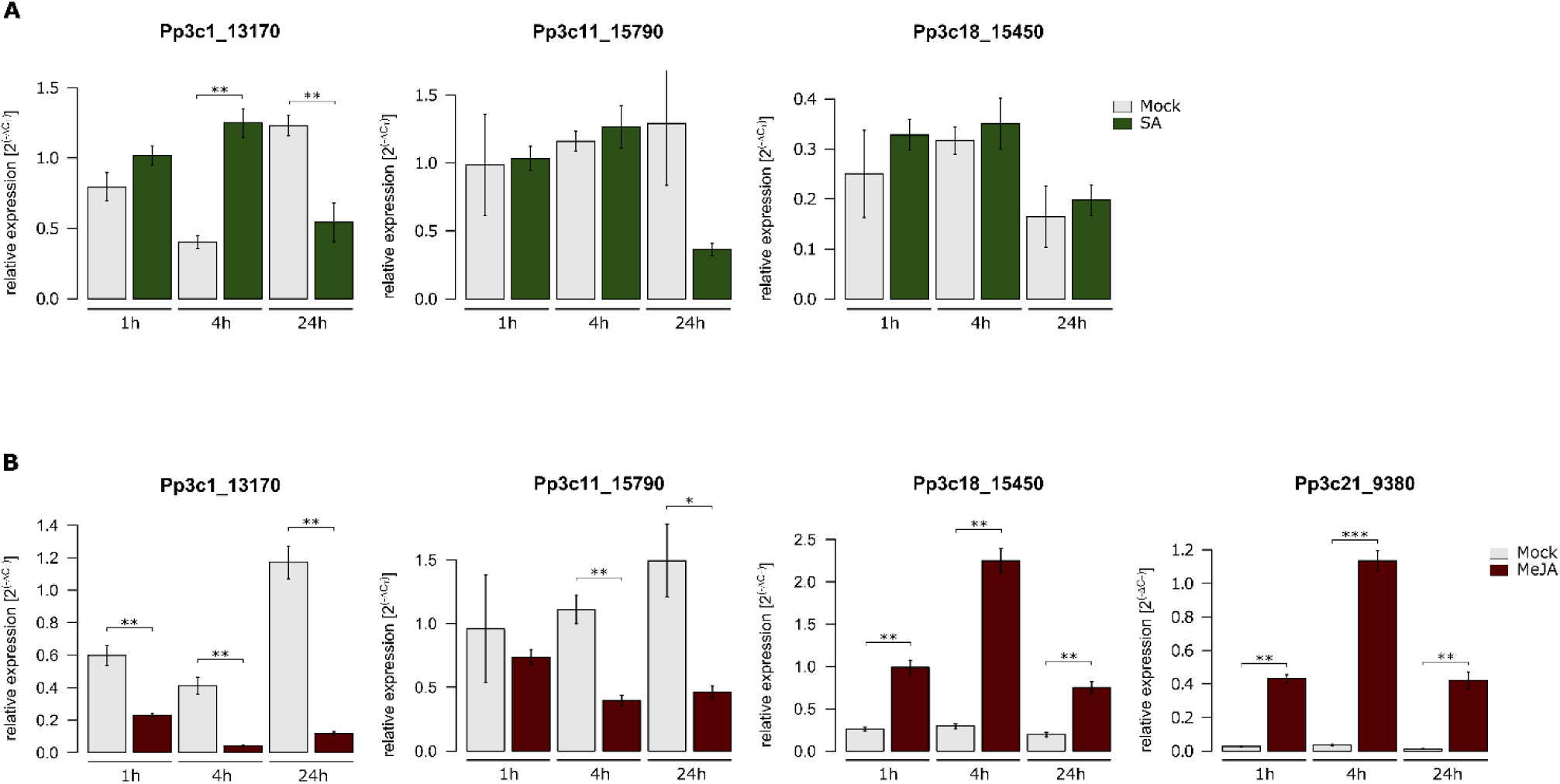
Physcomitrella *GAPDH*s gene expression upon phytohormone treatments. Protonema tissue was treated with 400 µM SA and 400 µM MeJA for 1h, 4h, and 24h. Upper panel: Treatment performed with 400 µM SA. Bottom panel: Treatment performed with 400 µM MeJA. Relative expression (2^(-ΔCT)^) is calculated against the reference genes *C45* (Pp3c9_13440) and *EF1α* (Pp3c27_2160) according to Livak & Schmittgen (2001). Mean values from three technical replicates with standard deviation are depicted. Statistical significance levels are based on a two-tailed Welsh t-test. *P* *>0.05, \*\**P* >0.005, \*\*\**P* >0.001.

According to this first analysis all selected candidate genes responded within 4 hours of treatment with MeJA, either being upregulated or downregulated (Figure 2). In the SA treatment the only response was also observed after 4 hours causing a significant upregulation of Pp3c1_13170. In consequence, we re-evaluated the response of the 4 candidates to a 4 hours MeJA treatment in biological replicates. Here, the trends in gene expression changes of all four candidate *GAPDH* genes were confirmed (Figure 3A). The two *GAPC* (Figure 3A, Pp3c18_15450, Pp3c21_9380) were significantly upregulated by 24- and 14-fold, respectively. Conversely, both genes *GAPA* genes (Figure 3A, Pp3c1_13170, Pp3c11_15790) were significantly downregulated by 6- and 14-fold respectively.

**Figure 3:**
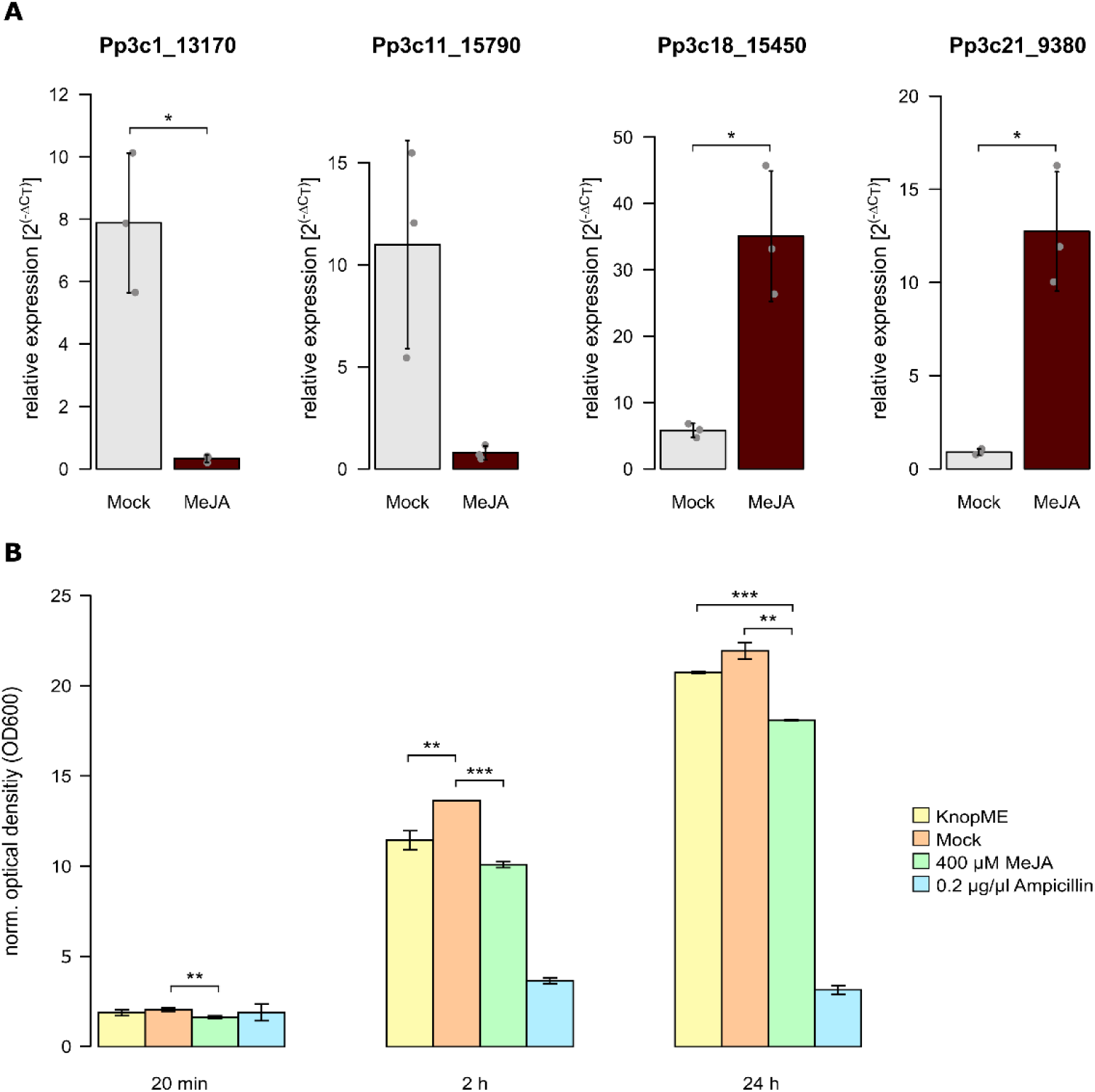
Expression analysis in biological triplicates and bioactivity assay of treated secretome samples A) Physcomitrella *GAPDH*s gene expression upon 4 hours of MeJA treatment in biological triplicates. Protonema tissue was treated with 400 µM MeJA for 4h. Relative expression (2^(-ΔCT)^) is calculated against the reference genes *C45* (Pp3c9_13440) and *LWD* (Pp3c22_18860) according to Livak & Schmittgen (2001). Mean values from three biological replicates (grey dots) with standard deviation are depicted. Values from each biological replicate are mean values from each three technical replicates. B) Bioactivity assay of secretome of MeJA-treated protonema suspension culture against *E. coli* K12. Antimicrobial activity was observed by measurement of optical density (A_600_) of bacterial cultures after 20 min, 2 hours and 24 hours incubation with secretome samples. KnopME: Physcomitrella culture medium, Mock: Culture Medium with addition of EtOH (solvent used for MeJA stock solutions). Mean values from three biological (A) and technical replicates (B) with standard deviation are depicted. Statistical significance levels were calculated with a two-tailed Welsh t-test. *P* *>0.05, \*\**P* >0.005, \*\*\**P* >0.001.

Since we observed significant changes in gene expression of the 4 *GAPDH*s in response to a treatment with a key hormone of the plant defense system, we were interested if within the applied timeframe an antimicrobial response could be triggered. To address this question, we performed a bioactivity assay with secretome samples of protonema treated for 4 hours with 400 µM MeJA. As antimicrobial compounds other than protein and peptides and the high salt concentration of the secretome samples might have an effect on bacterial growth, we lyophilized the secretome samples and subsequently employed C18-columns to remove salt and hydrophilic compounds as well to enrich proteins and peptides.

Since an antimicrobial activity against gram-negative bacteria was predicted for bioactive sequences in the selected GAPDHs (Table 1), we employed *E. coli* (K12) and monitored bacterial growth over time. Effectively, bacterial growth was significantly inhibited already 20 minutes after application of the treated secretome sample (Figure 3B). After 2 hours of incubation a growth inhibition of 26% was observed, which dropped to 17% after 24 hours of incubation. Consequently, the treatment of protonema with 400 µM MeJA for 4 hours triggers a remodeling of the secretome causing a bacteriostatic effect. Within the same time frame, the expression of cytosolic GAPDHs is upregulated while the expression of plastid GAPDHs is downregulated.

## Discussion

GAPDHs are known for their role in the glycolysis pathway and in the Calvin-Benson cycle but also for their magnitude of moonlight activities. In plants, GAPDHs were shown to play a role in abiotic stress response mechanisms, such as drought (Guo *et al*., 2012), salt (Lu *et al*., 2020), heat (Yang *et al*., 1993) and cold (Bae *et al*., 2003). GAPDHs were also shown to play an important role in pathogen response mechanisms. In this work we investigated the putative involvement of Physcomitrella GAPDHs in plant pathogen response by analyzing the inducibility of selected *GAPDH* genes upon SA- and MeJA-treatment, which allows to investigate the putative involvement of GAPDHs in the immune response against biotrophic and necrotrophic pathogens. SA is a positive regulator of immune responses against biotrophic pathogens, whereas JA is a positive regulator of immune responses against necrotrophic pathogens.

Our phylogenetical analysis resolved 17 phosphorylating GAPDHs from 4 different subgroups in the moss Physcomitrella. In contrast to other land plants analyzed in our study, Physcomitrella as well as *F. hygrometrica* have several additional *GAPDH* genes. For example, Arabidopsis has 2 *GAPA*s, 1 *GAPB*s, 2 *GAPC*s and 2 *GAPCp*s, while we identified 6 *GAPA*, 8 *GAPC* and 3 *GAPCp* genes in Physcomitrella. Physcomitrella, as well as *F. hygrometrica* and *S. angustifolium* do not possess any *GAPB* genes, confirming earlier studies showing that *GAPB* resulted from a gene duplication of *GAPA* at the beginning of land plant evolution (Petersen *et al.,* 2006; Martin & Cerff 2017). The additional 4 *PpGAPA*, 6 *PpGAPC* and 1 *PpGAPCp* genes might be a result of lineage-specific duplications. Physcomitrella was shown to have a higher abundance of proteins involved in metabolic processes due to genome duplication events (Rensing *et al*., 2007; Lang *et al*., 2005). Remarkably, the here analyzed Funariaceae have several more *GAPC* genes clustered in a separate subclade. Whether these additional *GAPC* genes produce functional proteins, needs to be further elucidated. The analyzed transcriptomic data showed that these genes are hardly expressed and no proteomic evidences for these GAPCs were found, further supporting the absence of any transcript and functional protein. Additionally, the GAPC Pp3c24_16410 was found to lack the catalytic cysteine. Several studies have shown that post-translational modification of the catalytic cysteine can play an important role in other non-glycolytic functions, such as apoptosis (reviewed in Nicholls *et al.,* 2012; Zaffagnini *et al.,* 2013). These results indicate that the once duplicated *GAPC* genes do not possess any function in Physcomitrella, and hence might represent pseudogenes. A large number of *GAPDH* pseudogenes were also described for other organisms, such as mouse (331 *GAPDH* pseudogenes) and rat (364 *GAPDH* pseudogenes) (Liu *et al*., 2009). These authors speculated that the high number of *GAPDH* genes might correlate with their moonlight activities.

We identified 10 GAPDHs across 10 mass spectrometry datasets and chose two photosynthetic GAPAs and two cytoplasmic glycolic GAPCs based on their proteomic and transcriptomic evidences and analyzed their expression upon MeJA- and SA-treatment. Remarkably, our results show contrasting gene regulation of the selected GAPDHs between the applied phytohormone treatments and within the same treatments we observed contrasting gene expression between the GAPDH subfamilies. We found that the analyzed *GAPA* (Pp3c11_15790) and the *GAPC* (Pp3c18_15450) were not responsive to SA treatments. Exclusively the *GAPA* Pp3c1_13170 displayed a significant short-term 3-fold upregulation after 4 hours of SA-treatment, before being significantly downregulated to its initial expression level after 24 hours of treatment.

In contrast, MeJA-treatment resulted in distinct expression profiles for the GAPDH-coding genes from different subfamilies. While the expression of both *GAPA*s was downregulated, a significant upregulation of both *GAPC*s was observed, with Pp3c21_9380 displaying an up to 33-fold upregulation after 4 hours and 24 hours of MeJA-treatment. The maintenance of increased gene expression over the course of 24 hours might indicate that the *GAPC* Pp3c21_9380 plays an important role in MeJA-mediated pathogen response mechanisms. Contrasting expression regulation of *GAPA/B* and *GAPC* genes was also observed after the induction of pathogen response of Arabidopsis with the elicitor flg22 (Henry *et al*., 2015). The authors describe a slight downregulation of the chloroplastic *GAPA*s and *GAPB*, and a two-fold upregulation of *GAPC1*, while *GAPC2* didn’t show any change in its gene expression. The authors hypothesized that the upregulation of the glycolytic GAPDH might be necessary to increase the ATP amounts as well as to increase the amount of ROS scavenging pyruvate.

The here shown contrasting expression of the GAPDHs in response to MeJA- and SA treatment might indicate a role of GAPDH and GAPDH-related antimicrobial peptides in the immune response against necrotrophic pathogens, which is positively regulated through the JA/ET pathway. While biotrophic pathogens feed on living cells, necrotrophic pathogens first induce host cell death to feed on the contents (Pieterse *et al*., 2012). The pathogen response mechanisms usually include the production of reactive oxygen species (ROS) and programmed cell death (hypersensitive response) at the infection site (Jones & Dangl, 2006). However, while ROS and apoptosis are powerful tools against biotrophic organisms, they are counterproductive for combatting necrotrophic pathogens. Several studies showed that GAPDHs can act as redox sensors and are involved in the inhibition of ROS production and in H_2_O_2_-mediated apoptosis (Baek *et al.,* 2008; Rius *et al.,* 2008; Yang & Zhai 2017; Henry *et al.,* 2015). Chloroplast-derived ROS play an important role in plant immune response (Littlejohn *et al*., 2021) and the downregulation of the chloroplastic *GAPA*s might be necessary to reduce the consumption of NADPH in the GAPA-catalyzed reaction, as NADPH is cofactor by many ROS scavenging enzymes, such as Glutathione Reductase (GR) which reduces GSSG to the antioxidant GSH (Das & Roychoudhury, 2014). Beside a putative indirect effect on pathogen response mechanisms, PpGAPDH might have a direct function in pathogen response *via* GAPDH-derived antimicrobial peptides. Several GAPDH-derived peptides from different organisms were shown to have antimicrobial properties (Branco *et al.,* 2018; Seo *et al.,* 2012) and a peptide derived from the PpGAPA Pp3c2_24160 showed antimicrobial activity (Fesenko *et al*., 2019). All here analyzed PpGAPDHs contain a sequence motif homologue to the antimicrobial peptides YFGAP and were predicted to have an antimicrobial activity against gram-negative bacteria. Plant antimicrobial peptides which are derived from functional precursor are generated by proteolytic processing (Tavormina *et al*., 2015). Nine of the 10 identified GAPDHs were identified in peptidome analyses and several GAPDH-derived peptides were exclusively found in phytohormone treated samples, further supporting the putative involvement of GAPDH-derived antimicrobial peptides in pathogen response in Physcomitrella. The direct function of the PpGAPCs-derived antimicrobial peptides as shown for GAPDHs from *T. albacares* (Seo *et al*., 2012), yeast (Branco *et al*., 2018) and a GAPA from Physcomitrella (Fesenko *et al*., 2019), is yet to be elucidated.

Our results identified two MeJA-inducible GAPDHs from Physcomitrella as putative positive regulators of the immune response against necrotrophic pathogens and negative regulators for immune responses against biotrophic pathogens.

## Material and Methods

### Phylogenetic analysis

We used the *Arabidopsis thaliana* GAPA protein sequence At1G12900 as the query in BLASTP-like searches with diamond (Buchfink *et al.,* 2021) against the proteomes of *Physcomitrium patens*, *Funaria hygrometrica, Sphagnum angustifolium, Marchantia polymorpha*, *Arabidopsis thaliana* and *Oryza sativa*. A multiple sequence alignment of the resulting proteins was performed with SEPP/UPP (Mirarab *et al.*, 2012; Nguyen *et al*., 2015) and trimmed with trimAl (Capella-Gutiérrez *et al.,* 2009) selecting a gap threshold of 0.1. Subsequently, we used RAxML (Stamatakis, 2014) to reconstruct a maximum-likelihood phylogenetic tree based on 1,000 rapid bootstrap replicates and automatic model selection.

### Computational predictions

Prediction of the sub cellular localization was performed using TargetP2.0 (Armenteros *et al*., 2019). To identify putative GAPDH-derived antimicrobial peptides we conducted a BLAST search using the DRAMP database (http://dramp.cpu-bioinfor.org; Shi *et al*., 2022). The prediction of the antimicrobial activity was performed using DRAMP3.0 prediction tools (Shi *et al*., 2022).

### Analysis of publicly available proteomic and transcriptomic datasets

We constructed a proteome database comprising mass spectrometry (MS) datasets of Physcomitrella proteome and peptidome analyses, which are publicly available. The following datasets were downloaded from the ProteomeXchange database (Deutsch *et al*., 2023; http://www.proteomexchange.org): PXD000227 (Fesenko *et al*., 2015), PXD009530 (Fesenko *et al*., 2019), PXD009532 (Fesenko *et al*., 2019), PXD002866 (Fesenko *et al*., 2016), PXD0012843 (Müller-Schüssele *et al*., 2020), PXD010964 (Filippova *et al*., 2019), PXD009517 (Hoernstein *et al*., 2018). Additionally, data published by Lehtonen *et al*. (2014) were used.

The MS data were reprocessed using Mascot Distiller V.2.7.0 (Matrix Science) using a database containing all V3.3 protein models of Physcomitrella (Lang *et al*., 2018) and an in-house contaminants database. Processing options were selected according to the MS instrument as well as the applied experimental setup described in the respective publications. For MS data generated with TripleTOF 5600 and Orbitrap Velos (PXD000227, PXD002866) instruments the fragment tolerance was set to 50 ppm and the parent mass tolerance was set to 20 ppm. For data generated with a QExactiveHF and QExactivePlus the fragment tolerance was set to

0.02 Da and the parent mass tolerance was set to 5.0 ppm. Carbamidomethylation (C, + 57.021464 Da) was set as fixed modification for all datasets except for the dataset PXD000227, were it was set as variable modification. The following variable modification were applied: pyro Glu (N-term Q, −17.026549 Da), hydroxylation (P, +15.9949 Da), oxidation (M, + 15.994915 Da), acetylation (Peptide N-term, + 42.010565 Da) and deamidation (N, + 0.984016 Da), sulfation (+ 79.96 Da). The latter one was not chosen for the dataset PXD000227. The enzyme specificity was set to tryptic (semi-specific free N-terminus) for the datasets (PXD002866, PXD0012843), with 2 possible miss cleavages. For peptidome analyses (PXD000227, PXD009530, PXD009532, PXD010964) the enzyme specificity was set to “non-specific”. Processing results were loaded into Scaffold^TM^ 4 (V4.11.1, Proteome Software) or Scaffold^TM^ 5 (V5.0.1, Proteome Software). Protein and peptide threshold were set to 99.9% and 95% respectively.

### Cultivation of Physcomitrella

Physcomitrella wild type (new botanical name: *Physcomitrium patens* (Hedw.) Mitt. (Medina *et al*., 2019), ecotype “Gransden 2004”was cultivated under axenic conditions in KnopME medium (Reski & Abel, 1985; (Knop medium pH 5.8: 1g/L Ca(NO_3_)_2_ x 4 H_2_O, 250 mg/L KCl, 250 mg/L KH_2_PO_4_, 250 mg/L MgSO_4_ x 7 H_2_O, 12.5 mg/L FeSO_4_ x 7 H_2_O; microelements (ME): 8.45 mg/L MnSO_4_ x 1 H_2_O, 4.31 mg/L ZnSO_4_ x 7 H_2_O, 3.09 mg/L H_3_BO_4_, 415 µg/L KJ, 121 µg/L Na_2_MoO_4_ x 2 H_2_O, 14.6 µg/L Co(NO_3_) x 6 H_2_O, 12.5 µg/L CuSO_4_ x 2 H_2_O). Moss protonema suspension culture was cultivated under standard long day conditions (16h light/8h dark photocycle; 55 µmol photons m^-2^ sec^-1^) at 23 °C. Protonema tissue was homogenized weekly with an ULTRA-TURRAX at 18000 rpm for 70 s. To verify the sterility of protonema suspension culture, small amounts of the homogenized protonema was given on LB-, TSA-2% Glucose-, and KnopME-medium agar plates (LB-agar pH 7.0: 10 g/L Tryptone, 5 g/L Yeast extract, 10 g/L NaCl, 15 g/L Agar; TSA-2% Glucose-Agar pH 7.5: 15 g/L Peptone from casein, 5 g/L Soya peptone, 5 g/L NaCl, 11g/L Glucose monohydrate, 15 g/L Agar) and checked for up to 6 weeks. The last homogenization process was performed three days prior to phytohormone treatment.

### Phytohormone treatment

Moss treatment with methyl jasmonate (MeJA) or salicylic acid (SA) was performed as followed: Moss protonema culture was adjusted to 50 mgDW/ml and was cultivated in KnopME either for 3 days for experiments in biological triplicates or were directly treated with phytohormones for experiments in technical replicates. The treatments were performed with 400 µM MeJA or 400 µM SA respectively. Phytohormone solutions were filtered *via* a 0.22 PES filter and were added slowly in drops. For the untreated samples the solvent used for the phytohormone solution (99.5% ethanol) was applied in an analogous manner. Samples were drawn after 1, 4 and 24 hours under sterile conditions and flash frozen in 100 mgFW portions. The harvested frozen plant material was disrupted in a bead mill for 3 min at 30 Hz using a 2 mL reaction tube with a glass and a tungsten carbide bead. The samples were stored at -80 °C.

### RNA isolation and reverse transcription to cDNA

RNA extraction was performed using TRIzol^TM^ (Invitrogen) according to the manufacturer’s instructions. The RNA concentration was determined photometrically (NanoDrop ND 1000, PEQLAB Biotechnologie GmbH, Erlangen). To check the RNA integrity and quality 500 ng RNA were analyzed *via* agarose gel electrophoresis. The residual DNA contained in the RNA samples was digested with DNase I (Thermo Fisher Scientific) for 1 hour at 37 °C followed by the inactivation of DNase I by the addition of EDTA (final conc.: 2.3 mM) and an incubation at 65 °C for 10 Min. The reverse transcription to cDNA was performed with the Reverse Transcription Reagents (Applied Biosystems, Waltham USA) using random hexamer primer. The cDNA synthesis was performed using the MultiScribe reverse transcriptase and the following synthesis steps: 25 °C for 10 min, 43 °C for 1h, and 95 °C for min. A non-transcribed control with no reverse transcriptase was performed to confirm a complete DNase I digestion.

Using the primary transcript coding sequences of the PpGAPDHs, we designed qPCR primer pairs (Supplementary Table 3) using the GeneScript “Real-time PCR (Taqman) Primer and Probes Tool” (https://www.genscript.com/tools/real-time-pcr-taqman-primer-design-tool/). The design was performed using the following characteristics: Amplicon size range of 50-100 nucleotides, a melting temperature of the primers of 60-68 °C with an optimum at 63 °C. The selection of primer pairs was performed based on their off-targets hits after BLAST search against the Physcomitrella V3.3 genome assembly (Lang *et al*., 2018). Primer were checked for their amplification efficiency as well as for possible off-targets. For this purpose, a 1:2 serial dilution of *P. patens* cDNA and a control without cDNA were used in a qPCR run as described below. Primer with an efficiency of 2 were chosen for further experiments. The absence of possible off-targets was ensured by analyzing the melting curves.

### Analyses of gene expression *via* qRT-PCR

Measurements were performed in technical replicates using 50 ng cDNA per triplicate. Reagents and each primer pair were checked for DNA contaminations *via* a control reaction with no template added.

Measurements were carried out in 96-well plates using the SensiFAST™ SYBR® Kit (Bioline, Luckenwalde, Germany) in a LightCycler® 480 (Roche). The qPCR run was performed as followed: First the samples were heated up to 95 °C for 2 min. For 45 consecutive cycles the plate was heated to 95 °C for 5 sec, cooled down to 60 °C for 10 sec, and subsequently heated up to 72 °C for 15 sec. After the 45 cycles the samples were brought to 65 °C, and gradually heated up to 95 °C. In the last step the samples were cooled down to 40 °C. During the cycles the samples were excited at 465 nm and the emission measured at λ=510 nm. For the relative quantification of expression levels of the GAPDHs, the two reference genes *EF1α* (Pp3c27_2160) and *C45* (Pp3c9_13440) were chosen for expression analysis at chosen time points in technical replicates. For the analyses using the *EF1α* as reference gene the efficiency was set to the value 1.7. For experiments were the *GAPDH*s expression levels were measured in biological triplicates the reference genes *C45* and *LWD* (Pp3c22_18860) were applied. The computational analysis of the expression levels was performed using the LightCycler® 480 software (Roche) and calculated relative to the reference genes according to (Livak & Schmittgen (2001). The relative expression levels are depicted as 2^(-ΔCT)^ (ΔCT = CT_[GOI]_ – CT_[reference]_). Primer pairs used in the experiment are listed in the Supplementary Table 3.

### Analysis of antimicrobial properties of secretome of MeJA treated samples

Moss protonema culture was treated with 400 µM MeJA for 4 hours as described above. Additionally, a Mock flask containing the same volume 99.5 % Ethanol, and a flask containing just the moss culture medium (KnopME) were prepared accordingly. The secretomes as well as the KnopME medium were lyophilized in 50 mL portions and stored at -20 °C. Three portions, corresponding to 150 mL culture, were pooled and desalted. For this procedure three portions of the lyophilized secretome were resuspended in 5 mL 0.1 % FA. This procedure and all the following ones were repeated for all sample types. SampliQ C18 ODS 100 mg columns (Agilent Technologies, Santa Clara, USA) were prepared as followed: For the activation of the membrane, 3 mL 100 % ACN were used to wash the column by pushing the liquid through the column with a syringe. The column was subsequently washed with 3 mL 0.1 % FA before adding the resuspended sample and pushed through the column using a syringe. The column was then washed twice with 0.1 % FA before eluting the peptides in a fresh protein low binding tube with 3 mL 80 % ACN, 0.1 % FA. The samples were dried in a vacuum concentrator and resuspended in 2 mL LB medium, and filtered through a 0.22 µM PES filter. The filter was washed with 1 mL LB medium and added to the filtered sample and imminently used for the treatment of the bacterial culture, which was prepared as followed: *E. coli* K12 were cryo conserved in 10 % DMSO until use. The bacterial culture was grown in LB medium over night at 37 °C and subsequently 100 µL each were mixed with 1.9 mL LB medium and 1 mL of the filtered secretome sample. This step was repeated for all sample types. For the positive control ampicillin was added (0.2 µg/µL final conc.) instead of the sample to the bacterial culture. The bacterial culture was subsequently cultivated at 37 °C, samples drawn at different time points (t_0_=0 min, 20 min, 2 h, and 24 h) and the bacterial growth monitored by measuring the optical density photometrically (A_600_). LB medium was used as blank and the samples drawn at t_0_ was used to normalize the corresponding sample.

## Statistical analyses

Statistical differences were analyzed *via* a two-tailed Welsh’s t-test (unequal variances) using Microsoft Excel (Microsoft Corporation, 2018. *Microsoft Excel*, https://office.microsoft.com/excel) and Graphpad (https://www.graphpad.com/quickcalcs/ttest2/).

## Declaration

The authors declare no competing interests.

## Author Contributions

A.A.M. designed the study, performed experiments, analyzed data, and wrote the manuscript. S.N.W.H. analyzed data and helped writing the manuscript. N.v.G. analyzed data and helped writing the manuscript. J.O.P. performed experiments. R.R. designed and supervised the study, acquired funding, and wrote the manuscript. All authors approved the final version of the manuscript.

## Acknowledgements

This work was supported by the German Federal Ministry of Education and Research (BMBF) Project ID 01DK20064 and by the German Research Foundation (DFG) under Germany’s Excellence Strategy (CIBSS – EXC-2189 – Project ID 390939984).

## Supplementary Data

**Supplementary Figure 1:**
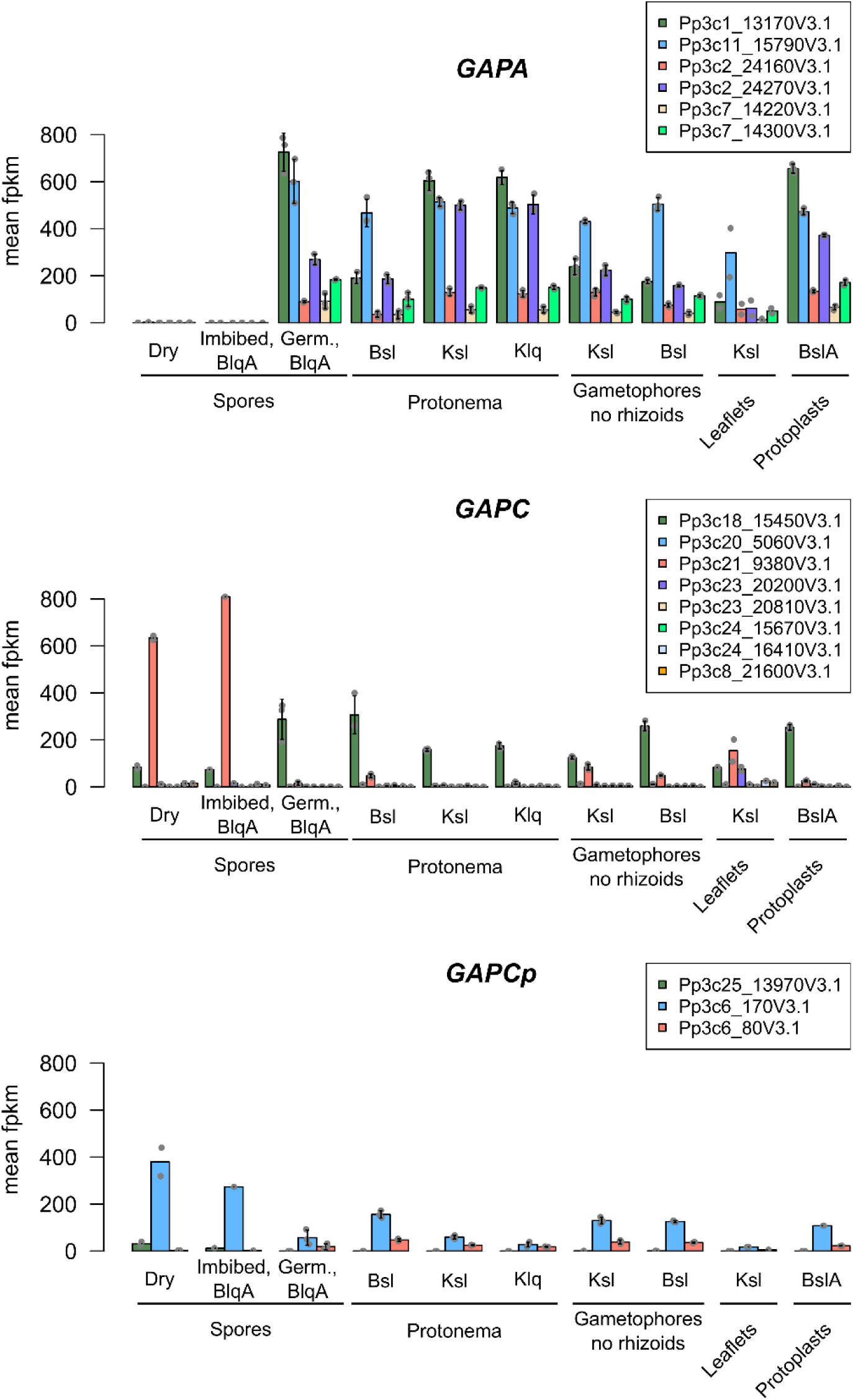
Expression levels of Physcomitrella *GAPA*, *GAPC*, and *GAPCp* genes in various developmental stages and cell culture conditions. Data were downloaded from PEATmoss (PEATmoss; Fernandez-Pozo *et al*., 2020; https://peatmoss.plantcode.cup.uni-freiburg.de). FPKM (fragments per kilobase million) values and standard deviation are depicted. Data points represent biological duplets or triplicates. Abbreviations: B: BCD medium, Germ.: germinating, lq: liquid, A: ammonium tartrate, sl: solid, K: Knop medium.

**Supplementary Figure 2:**
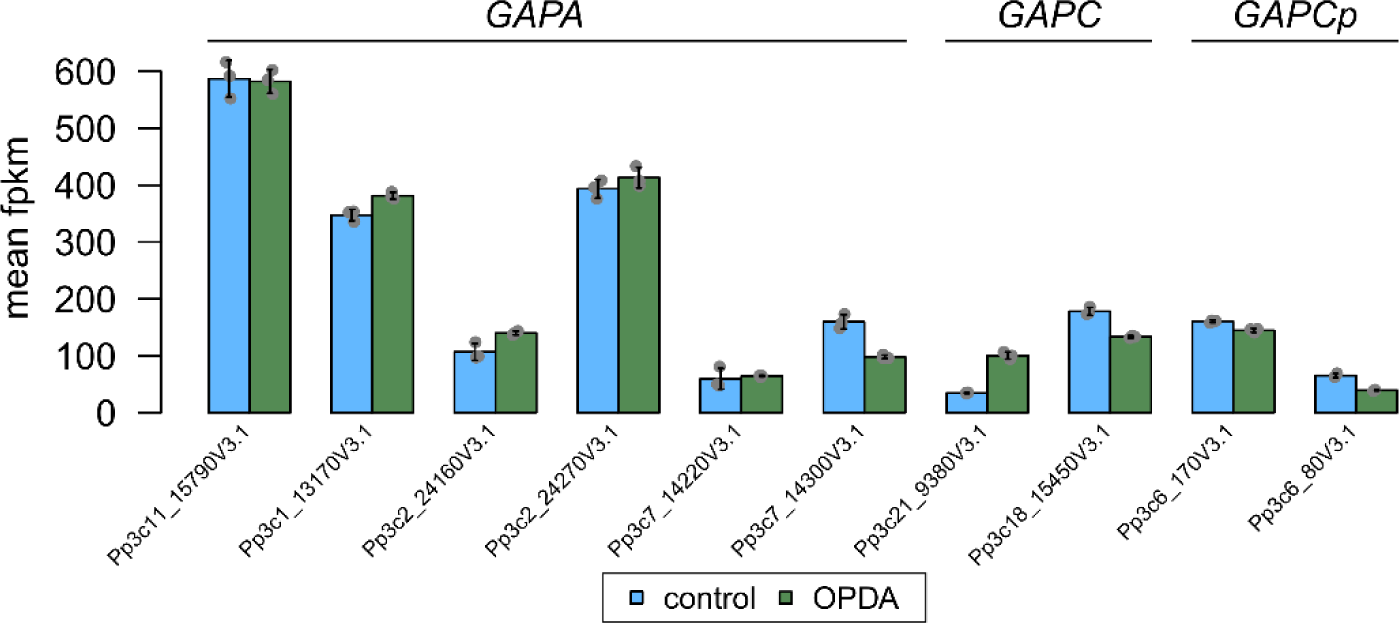
Expression of *GAPA*, *GAPC*, and *GAPCp* genes upon OPDA treatment. Data were downloaded from PEATmoss (Fernandez-Pozo *et al*., 2020; https://peatmoss.plantcode.cup.uni-freiburg.de). FPKM (fragments per kilobase million) values and standard deviation are depicted. Data points represent biological triplicates.

**Supplementary Table 1:** Subcellular localization of Physcomitrella GAPDHs. Prediction was performed using TargetP2.0 (Armenteros *et al*., 2019). Abbreviations: noTP: no transit peptide, SP: Signal peptide, mTP: mitochondrial transit peptide, cTP: chloroplast transit peptide, luTP: thylakoid luminal transit peptide, CS: cleavage site.

| Acc. ID | GAPDH subfamily | Prediction | noTP | SP | mTP | cTP | luTP | CS Position |
| --- | --- | --- | --- | --- | --- | --- | --- | --- |
| Pp3c1_13170V3.1 | GAPA | cTP | 0.000459 | 0.000006 | 0.000036 | 0.932516 | 0.066983 | CS pos: 69-70. TEA-KI. Pr: 0.2957 |
| Pp3c1_13170V3.2 | GAPA | cTP | 0.000576 | 0.00001 | 0.000039 | 0.920303 | 0.079073 | CS pos: 69-70. TEA-KI. Pr: 0.2968 |
| Pp3c11_15790V3.1 | GAPA | cTP | 0.000624 | 0.000002 | 0.000085 | 0.891688 | 0.107601 | CS pos: 69-70. TEA-KI. Pr: 0.2613 |
| Pp3c11_15790V3.3 | GAPA | noTP | 0.92581 | 0.002404 | 0.041951 | 0.029603 | 0.000232 |  |
| Pp3c11_15790V3.4 | GAPA | noTP | 0.999967 | 0.000031 | 0.000002 | 0 | 0 |  |
| Pp3c11_15790V3.5 | GAPA | cTP | 0.002599 | 0.000012 | 0.000112 | 0.936757 | 0.06052 | CS pos: 69-70. TEA-KI. Pr: 0.2612 |
| Pp3c11_15790V3.6 | GAPA | cTP | 0.002226 | 0.000007 | 0.000093 | 0.935558 | 0.062115 | CS pos: 69-70. TEA-KI. Pr: 0.2610 |
| Pp3c11_15790V3.7 | GAPA | cTP | 0.003296 | 0.000003 | 0.000165 | 0.938603 | 0.057933 | CS pos: 69-70. TEA-KI. Pr: 0.2712 |
| Pp3c2_24160V3.1 | GAPA | cTP | 0.003994 | 0.000012 | 0.000258 | 0.923884 | 0.071852 | CS pos: 62-63. TEA-KI. Pr: 0.2587 |
| Pp3c2_24270V3.2 | GAPA | noTP | 0.696543 | 0.121723 | 0.105407 | 0.069457 | 0.006871 |  |
| Pp3c2_24270V3.1 | GAPA | cTP | 0.003248 | 0.000016 | 0.000172 | 0.948237 | 0.048326 | CS pos: 69-70. TEA-KI. Pr: 0.2391 |
| Pp3c2_24270V3.3 | GAPA | noTP | 0.999844 | 0.000152 | 0.000004 | 0 | 0 |  |
| Pp3c7_14220V3.1 | GAPA | cTP | 0.001224 | 0.000018 | 0.000089 | 0.963499 | 0.03517 | CS pos: 69-70. TEA-KI. Pr: 0.3045 |
| Pp3c7_14300V3.1 | GAPA | cTP | 0.087366 | 0.052147 | 0.187376 | 0.502411 | 0.1707 | CS pos: 63-64. TEA-KI. Pr: 0.3878 |
| Pp3c7_14300V3.3 | GAPA | mTP | 0.107917 | 0.019418 | 0.62469 | 0.235005 | 0.012969 | CS pos: 83-84. SRA-VT. Pr: 0.0776 |
| Pp3c18_15450V3.1 | GAPC | noTP | 0.984573 | 0.012082 | 0.003017 | 0.00029 | 0.000038 |  |
| Pp3c20_5060V3.1 | GAPC | noTP | 0.989739 | 0.009236 | 0.000496 | 0.000524 | 0.000005 |  |
| Pp3c20_5060V3.2 | GAPC | noTP | 0.998113 | 0.001552 | 0.000297 | 0.000037 | 0.000001 |  |
| Pp3c20_5060V3.5 | GAPC | noTP | 0.99861 | 0.000802 | 0.000451 | 0.000132 | 0.000005 |  |
| Pp3c20_5060V3.6 | GAPC | noTP | 0.998113 | 0.001552 | 0.000297 | 0.000037 | 0.000001 |  |
| Pp3c21_9380V3.1 | GAPC | noTP | 0.985349 | 0.011756 | 0.002566 | 0.0003 | 0.000029 |  |
| Pp3c23_20200V3.1 | GAPC | noTP | 0.994349 | 0.00044 | 0.005119 | 0.000068 | 0.000023 |  |
| Pp3c23_20810V3.1 | GAPC | noTP | 0.983072 | 0.00137 | 0.010025 | 0.005412 | 0.00012 |  |
| Pp3c23_20810V3.6 | GAPC | noTP | 0.772175 | 0.111676 | 0.115041 | 0.00087 | 0.000237 |  |
| Pp3c24_15670V3.1 | GAPC | noTP | 0.996287 | 0.00156 | 0.001487 | 0.00064 | 0.000025 |  |
| Pp3c24_16410V3.1 | GAPC | noTP | 0.998784 | 0.000712 | 0.000466 | 0.000035 | 0.000003 |  |
| Pp3c24_16410V3.2 | GAPC | noTP | 0.998784 | 0.000712 | 0.000466 | 0.000035 | 0.000003 |  |
| Pp3c24_16410V3.3 | GAPC | noTP | 0.99975 | 0.000198 | 0.000052 | 0 | 0 |  |
| Pp3c24_16410V3.4 | GAPC | noTP | 0.99975 | 0.000198 | 0.000052 | 0 | 0 |  |
| Pp3c24_16410V3.6 | GAPC | noTP | 0.987533 | 0.001883 | 0.005188 | 0.005349 | 0.000047 |  |
| Pp3c8_21600V3.1 | GAPC | noTP | 0.997852 | 0.001562 | 0.000558 | 0.000023 | 0.000005 |  |
| Pp3c25_13970V3.1 | GAPCp | SP | 0.093419 | 0.447644 | 0.253959 | 0.182914 | 0.022064 | CS pos: 23-24. AQA-HF. Pr: 0.8379 |
| Pp3c6_170V3.1 | GAPCp | cTP | 0.002986 | 0.000007 | 0.000345 | 0.996631 | 0.000031 | CS pos: 75-76. TTR-AT. Pr: 0.3809 |
| Pp3c6_170V3.4 | GAPCp | noTP | 0.612193 | 0.000078 | 0.004236 | 0.383347 | 0.000146 |  |
| Pp3c6_80V3.1 | GAPCp | cTP | 0.00024 | 0.000006 | 0.093763 | 0.905476 | 0.000514 | CS pos: 76-77. VRA-TA. Pr: 0.6392 |

**Supplementary Table 2:** List of ProteomeXchange (http://www.proteomexchange.org; Deutsch *et al*., 2023) datasets identifiers used to construct proteome databank.

| ProteomXchange ID | Reference | Phytohormone treatment | Peptidome or proteome analyses | Plant tissue or sample type |
| --- | --- | --- | --- | --- |
| PXD010964 | Filippova *et al.*, 2019 | SA | Peptidome | Secretome |
| PXD009532 | Fesenko *et al.*, 2019 | MeJA and SA | Peptidome | Protonema |
| PXD009530 | Fesenko *et al.*, 2019 | MeJA | Peptidome | Secretome |
| PXD000227 | Fesenko *et al.*, 2015 | - | Peptidome | Gametophores, Protonema, Protoplasts |
| PXD002866 | Fesenko *et al.*, 2016 | - | Proteome | Protonema, Protoplasts |
| PXD0012843 | Müller-Schüssele *et al.*, 2019 | - | Proteome | Protonema |
| PXD009517 | Hoernstein *et al.*, 2018 | - | Proteome | Secretome |
| - | Lehtonen *et al.*, 2014 | Chitosan | Proteome | Secretome |

**Supplementary Table 3:**
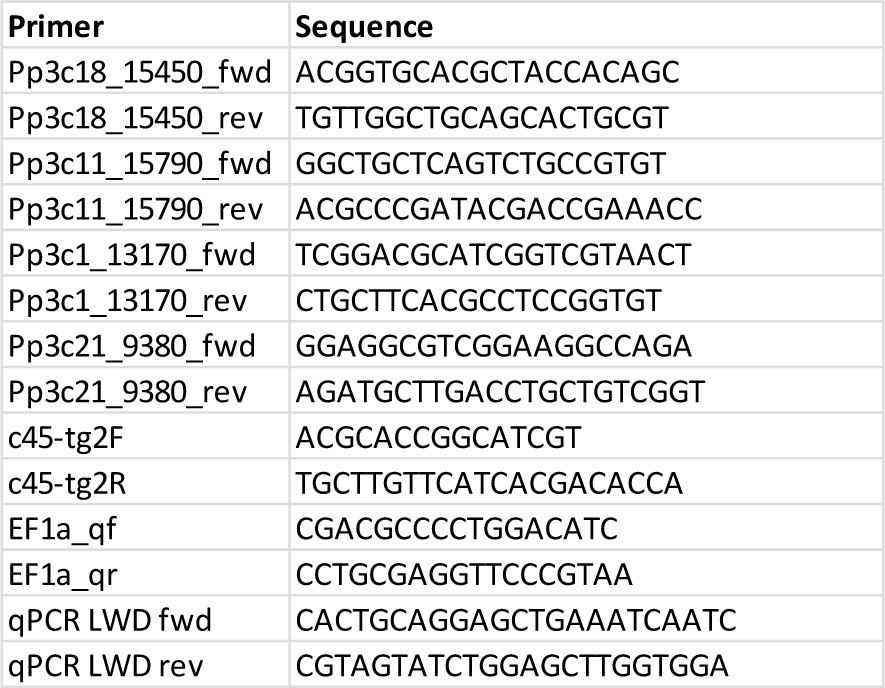
Primer pairs used for qRT-PCR analyses of target genes.

## References

1. Arif, M.A., Hiss, M., Tomek, M., Busch, H., Meyberg, R., Tintelnot, S., Reski, R., Rensing, S.A. & Frank, W. ABA-induced vegetative diaspore formation in *Physcomitrella patens*. Front. Plant Sci. 10, 315 (2019).

2. Armenteros, J.J.A., Salvatore, M., Emanuelsson, O., Winther, O., Von Heijne, G., Elofsson, A. & Nielsen, H. Detecting sequence signals in targeting peptides using deep learning. Life Sci. Alliance 2, e201900429 (2019).

3. Bae, M.S., Cho, E.J., Choi, E.Y. & Park, O.K. Analysis of the Arabidopsis nuclear proteome and its response to cold stress. Plant J. 36, 652–663 (2003).

4. Baek, D., Jin, Y., Jeong, J.C., Lee, H.J., Moon, H., Lee, J., Shin, D., Kang, C.H., Kim, D.H., Nam, J., Lee, S.Y. & Yun, D.J. Suppression of reactive oxygen species by glyceraldehyde-3-phosphate dehydrogenase. Phytochemistry 69, 333–338 (2008).

5. Bari, R. & Jones, J.D.G. Role of plant hormones in plant defence responses. Plant Mol. Biol. 69, 473–488 (2009).

6. Bennett, T.A., Liu, M.M., Aoyama, T., Bierfreund, N.M., Braun, M., Coudert, Y., Dennis, R.J., O’Connor, D., Wang, X.Y., White, C.D., Decker, E.L., Reski, R. & Harrison, C.J. Plasma membrane-targeted PIN proteins drive shoot development in a moss. Curr. Biol. 24, 2776–2785 (2014).

7. Bowman, J.L., Kohchi, T., Yamato, K.T., Jenkins, J., Shu, S., Ishizaki, K., Yamaoka, S., Nishihama, R., Nakamura, Y., Berger, F., Adam, C., Aki, S.S., Althoff, F., Araki, T., Arteaga-Vazquez, M.A., et al. Insights into land plant evolution garnered from the *Marchantia polymorpha* Genome. Cell 171, 287–304 (2017).

8. Branco, P., Albergaria, H., Arneborg, N. & Prista, C. Effect of GAPDH-derived antimicrobial peptides on sensitive yeasts cells: membrane permeability, intracellular pH and H + -influx/-efflux rates. FEMS Yeast Res. 18, foy030 (2018).

9. Buchfink, B., Reuter, K. & Drost, H.G. Sensitive protein alignments at tree-of-life scale using DIAMOND. Nat. Methods 18, 366–368 (2021).

10. Butterfield, D.A., Hardas, S.S. & Bader Lange, M.L. Oxidatively modified Glyceraldehyde-3-Phosphate Dehydrogenase (GAPDH) and Alzheimer disease: Many pathways to neurodegeneration. J. Alzheimer’s Dis. 20, 369–393 (2010).

11. Capella-Gutiérrez, S., Silla-Martínez, J.M. & Gabaldón, T. trimAl: A tool for automated alignment trimming in large-scale phylogenetic analyses. Bioinformatics 25, 1972–1973 (2009).

12. Cheng, C.Y., Krishnakumar, V., Chan, A.P., Thibaud-Nissen, F., Schobel, S. & Town, C.D. Araport11: a complete reannotation of the *Arabidopsis thaliana* reference genome. Plant J. 89, 789–804 (2017).

13. Colell, A., Ricci, J.-E., Tait, S., Milasta, S., Maurer, U., Bouchier-Hayes, L., Fitzgerald, P., Guio-Carrion, A., Waterhouse, N.J., Li, C.W., Mari, B., Barbry, P., Newmeyer, D.D., Beere, H.M. & Green, D.R. GAPDH and autophagy preserve survival after apoptotic Cytochrome c release in the absence of Caspase activation. Cell 129, 983–997 (2007).

14. Colell, A., Green, D.R. & Ricci, J.-E. Novel roles for GAPDH in cell death and carcinogenesis. Cell Death Differ. 16, 1573–1581 (2009).

15. Das, K. & Roychoudhury, A. Reactive oxygen species (ROS) and response of antioxidants as ROS-scavengers during environmental stress in plants. *Front*. Environ. Sci. 2, 53 (2014).

16. Decker, E.L., Alder, A., Hunn, S., Ferguson, J., Lehtonen, M.T., Scheler, B., Kerres, K.L., Wiedemann, G., Safavi-Rizi, V., Nordzieke, S., Balakrishna, A., Baz, L., Avalos, J., Valkonen, J.P.T., Reski, R., et al. Strigolactone biosynthesis is evolutionarily conserved, regulated by phosphate starvation and contributes to resistance against phytopathogenic fungi in a moss, *Physcomitrella patens*. New Phytol. 216, 455–468 (2017).

17. Deutsch, E.W., Bandeira, N., Perez-Riverol, Y., Sharma, V., Carver, J.J., Mendoza, L., Kundu, D.J., Wang, S., Bandla, C., Kamatchinathan, S., Hewapathirana, S., Pullman, B.S., Wertz, J., Sun, Z., Kawano, S., et al. The ProteomeXchange consortium at 10 years: 2023 update. Nucleic Acids Res. 51, D1539–D1548 (2023).

18. Fernandez-Pozo, N., Haas, F.B., Meyberg, R., Ullrich, K.K., Hiss, M., Perroud, P.F., Hanke, S., Kratz, V., Powell, A.F., Vesty, E.F., Daum, C.G., Zane, M., Lipzen, A., Sreedasyam, A., Grimwood, J., et al. PEATmoss (Physcomitrella Expression Atlas Tool): a unified gene expression atlas for the model plant *Physcomitrella patens*. Plant J. 102, 165–177 (2020).

19. Fesenko, I.A., Arapidi, G.P., Skripnikov, A.Y., Alexeev, D.G., Kostryukova, E.S., Manolov, A.I., Altukhov, I.A., Khazigaleeva, R.A., Seredina, A. V., Kovalchuk, S.I., Ziganshin, R.H., Zgoda, V.G., Novikova, S.E., Semashko, T.A., Slizhikova, D.K., et al. Specific pools of endogenous peptides are present in gametophore, protonema, and protoplast cells of the moss *Physcomitrella patens*. BMC Plant Biol. 15, 1–16 (2015).

20. Fesenko, I., Seredina, A., Arapidi, G., Ptushenko, V., Urban, A., Butenko, I., Kovalchuk, S., Babalyan, K., Knyazev, A., Khazigaleeva, R., Pushkova, E., Anikanov, N., Ivanov, V. & Govorun, V.M. The *Physcomitrella patens* chloroplast proteome changes in response to protoplastation. Front. Plant Sci. 7, 1661 (2016).

21. Fesenko, I., Azarkina, R., Kirov, I., Kniazev, A., Filippova, A., Grafskaia, E., Lazarev, V., Zgoda, V., Butenko, I., Bukato, O., Lyapina, I., Nazarenko, D., Elansky, S., Mamaeva, A., Ivanov, V., et al. Phytohormone treatment induces generation of cryptic peptides with antimicrobial activity in the moss *Physcomitrella patens*. BMC Plant Biol. 19, 1–16 (2019).

22. Filippova, A., Lyapina, I., Kirov, I., Zgoda, V., Belogurov, A., Kudriaeva, A., Ivanov, V. & Fesenko, I. Salicylic acid influences the protease activity and posttranslation modifications of the secreted peptides in the moss *Physcomitrella patens*. J. Pept. Sci. 25, e3138 (2019).

23. Goodstein, D.M., Shu, S., Howson, R., Neupane, R., Hayes, R.D., Fazo, J., Mitros, T., Dirks, W., Hellsten, U., Putnam, N. & Rokhsar, D.S. Phytozome: A comparative platform for green plant genomics. Nucleic Acids Res. 40, D1178–D1186 (2012).

24. Guillory, A. & Bonhomme, S. Phytohormone biosynthesis and signaling pathways of mosses. Plant Mol. Biol. 107, 245–277 (2021).

25. Guo, L., Devaiah, S.P., Narasimhan, R., Pan, X., Zhang, Y., Zhang, W. & Wang, X. Cytosolic Glyceraldehyde-3-Phosphate Dehydrogenases interact with Phospholipase Dδ to transduce hydrogen peroxide signals in the Arabidopsis response to stress. Plant Cell 24, 2200–2212 (2012).

26. Habenicht, A., Hellman, U. & Cerff, R. Non-phosphorylating GAPDH of higher plants is a member of the Aldehyde Dehydrogenase superfamily with no sequence homology to phosphorylating GAPDH. J. Mol. Biol. 237, 165–171 (1994)

27. Hara, M.R., Agrawal, N., Kim, S.F., Cascio, M.B., Fujimuro, M., Ozeki, Y., Takahashi, M., Cheah, J.H., Tankou, S.K., Hester, L.D., Ferris, C.D., Hayward, D.S., Snyder, S.H. & Sawa, A. S-nitrosylated GAPDH initiates apoptotic cell death by nuclear translocation following Siah1 binding. Nat. Cell Biol. 7, 665–674 (2005).

28. Healey, A.L., Piatkowski, B., Lovell, J.T., Sreedasyam, A., Carey, S.B., Mamidi, S., Shu, S., Plott, C., Jenkins, J., Lawrence, T., Aguero, B., Carrell, A.A., Nieto-Lugilde, M., Talag, J., Duffy, A., et al. Newly identified sex chromosomes in the *Sphagnum* (peat moss) genome alter carbon sequestration and ecosystem dynamics. Nat. Plants 9, 238–254 (2023).

29. Henry, E., Fung, N., Liu, J., Drakakaki, G. & Coaker, G. Beyond Glycolysis: GAPDHs are multi-functional enzymes involved in regulation of ROS, autophagy, and plant immune responses. PLoS Genet. 11, e1005199 (2015).

30. Hildebrandt, T., Knuesting, J., Berndt, C., Morgan, B. & Scheibe, R. Cytosolic thiol switches regulating basic cellular functions: GAPDH as an information hub? Biol. Chem. 396, 523–537 (2015).

31. Hoernstein, S.N.W., Fode, B., Wiedemann, G., Lang, D., Niederkrüger, H., Berg, B., Schaaf, A., Frischmuth, T., Schlosser, A., Decker, E.L. & Reski, R. Host cell proteome of *Physcomitrella patens* harbors proteases and protease inhibitors under bioproduction conditions. J. Proteome Res. 17, 3749–3760 (2018).

32. Itakura, M., Kubo, T., Kaneshige, A. & Nakajima, H. Glyceraldehyde-3-phosphate dehydrogenase regulates activation of c-Jun N-terminal kinase under oxidative stress. Biochem. Biophys. Res. Commun. 657, 1–7 (2023).

33. Jones, J.D.G. & Dangl, J.L. The plant immune system. Nature 444, 323–329 (2006).

34. Kim, S.C., Guo, L. & Wang, X. Nuclear moonlighting of cytosolic glyceraldehyde-3-phosphate dehydrogenase regulates Arabidopsis response to heat stress. Nat. Commun. 11, 1–15 (2020).

35. Kirbis, A., Waller, M., Ricca, M., Bont, Z., Neubauer, A., Goffinet, B. & Szövényi, P. Transcriptional landscapes of divergent sporophyte development in two mosses, *Physcomitrium (Physcomitrella) patens* and *Funaria hygrometrica*. Front. Plant Sci. 11, 747 (2020).

36. Lang, D., Eisinger, J., Reski, R. & Rensing, S.A. Representation and high-quality annotation of the *Physcomitrella patens* transcriptome demonstrates a high proportion of proteins involved in metabolism in mosses. Plant Biol. 7, 238–250 (2005).

37. Lang, D., Ullrich, K.K., Murat, F., Fuchs, J., Jenkins, J., Haas, F.B., Piednoel, M., Gundlach, H., Van Bel, M., Meyberg, R., Vives, C., Morata, J., Symeonidi, A., Hiss, M., Muchero, W., et al. The *Physcomitrella patens* chromosome-scale assembly reveals moss genome structure and evolution. Plant J. 93, 515– 533 (2018).

38. Laxalt, A.M., Cassial, R.O., Sanllorenti, P.M., Madrid, E.A., Andreu, A.B., Raû, G., Conde, R.D. & Lamattina, L. Accumulation of cytosolic glyceraldehyde-3-phosphate dehydrogenase RNA under biological stress conditions and elicitor treatments in potato. Plant Mol. Biol. 30, 961–972 (1996).

39. Lehtonen, M.T., Takikawa, Y., Rönnholm, G., Akita, M., Kalkkinen, N., Ahola-Iivarinen, E., Somervuo, P., Varjosalo, M. & Valkonen, J.P.T. Protein secretome of moss plants (*Physcomitrella patens*) with emphasis on changes induced by a fungal elicitor. J. Proteome Res. 13, 447–459 (2014).

40. Li, X., Wei, W., Li, F., Zhang, L., Deng, X., Liu, Y. & Yang, S. The plastidial Glyceraldehyde-3-Phosphate Dehydrogenase is critical for abiotic stress response in wheat. Int. J. Mol. Sci. 20, Article 1104 (2019).

41. Lindner, A.C., Lang, D., Seifert, M., Podlešáková, K., Novák, O., Strnad, M., Reski, R. & von Schwartzenberg, K. Isopentenyltransferase-1 (IPT1) knockout in Physcomitrella together with phylogenetic analyses of IPTs provide insights into evolution of plant cytokinin biosynthesis. J. Exp. Bot. 65, 2533–2543 (2014).

42. Littlejohn, G.R., Breen, S., Smirnoff, N. & Grant, M. Chloroplast immunity illuminated. New Phytol. 229, 3088–3107 (2021).

43. Liu, Y.-J., Zheng, D., Balasubramanian, S., Carriero, N., Khurana, E., Robilotto, R. & Gerstein, M.B. Comprehensive analysis of the pseudogenes of glycolytic enzymes in vertebrates: the anomalously high number of GAPDH pseudogenes highlights a recent burst of retrotrans-positional activity. BMC Genomics 10, 480 (2009).

44. Livak, K.J. & Schmittgen, T.D. Analysis of relative gene expression data using real-time quantitative PCR and the 2^-ΔΔCT^ method. Methods 25, 402–408 (2001).

45. Lu, Q., Meng, X., Yang, F., Liu, X. & Cui, J. Characterization of LcGAPC and its transcriptional response to salt and alkali stress in two ecotypes of *Leymus chinensis* (Trin.) Tzvelev. Biotechnol. Biotechnol. Equip. 34, 115–125 (2020).

46. Lüth, V.M., Rempfer, C., van Gessel, N., Herzog, O., Hanser, M., Braun, M., Decker, E.L. & Reski, R. A Physcomitrella PIN protein acts in spermatogenesis and sporophyte retention. New Phytol. 237, 2118– 2135 (2023).

47. Martin, W.F. & Cerff, R. Physiology, phylogeny, early evolution, and GAPDH. Protoplasma 254, 1823–1834 (2017).

48. Medina, R., Johnson, M.G., Liu, Y., Wickett, N.J., Shaw, A.J. & Goffinet, B. Phylogenomic delineation of *Physcomitrium* (Bryophyta: Funariaceae) based on targeted sequencing of nuclear exons and their flanking regions rejects the retention of *Physcomitrella*, *Physcomitridium* and *Aphanorrhegma*. J. Syst. Evol. 57, 404–417 (2019).

49. Mirarab, S., Nguyen, N. & Warnow, T. SEPP: SATé-enabled phylogenetic placement. Biocomputing 2012, 247–258 (2012).

50. Müller-Schüssele, S.J., Wang, R., Gütle, D.D., Romer, J., Rodriguez-Franco, M., Scholz, M., Buchert, F., Lüth, V.M., Kopriva, S., Dörmann, P., Schwarzländer, M., Reski, R., Hippler, M. & Meyer, A.J. Chloroplasts require glutathione reductase to balance reactive oxygen species and maintain efficient photosynthesis. Plant J. 103, 1140–1154 (2020).

51. Muthamilarasan, M. & Prasad, M. Plant innate immunity: An updated insight into defense mechanism. J. Biosci. 38, 433–449 (2013).

52. Nguyen, N.P.D., Mirarab, S., Kumar, K. & Warnow, T. Ultra-large alignments using phylogeny-aware profiles. Genome Biol. 16, 1–15 (2015).

53. Nicholls, C., Li, H. & Liu, J.-P. GAPDH: A common enzyme with uncommon functions. Clin. Exp. Pharmacol. Physiol. 39, 674–679 (2012).

54. Ouyang, S., Zhu, W., Hamilton, J., Lin, H., Campbell, M., Childs, K., Thibaud-Nissen, F., Malek, R.L., Lee, Y., Zheng, L., Orvis, J., Haas, B., Wortman, J. & Buell, R.C. The TIGR rice genome annotation resource: Improvements and new features. Nucl. Acids Res. 35, D883–D887 (2007).

55. Petersen, J., Brinkmann, H. & Cerff, R. Origin, evolution, and metabolic role of a novel glycolytic GAPDH enzyme recruited by land plant plastids. J. Mol. Evol. 57, 16–26 (2003).

56. Petersen, J., Teich, R., Becker, B., Cerff, R. & Brinkmann, H. The GapA/B gene duplication marks the origin of Streptophyta (Charophytes and Land Plants). Mol. Biol. Evol. 23, 1109–1118 (2006).

57. Pieterse, C.M.J., Van der Does, D., Zamioudis, C., Leon-Reyes, A. & Van Wees, S.C.M. Hormonal modulation of plant immunity. Annu. Rev. Cell Dev. Biol. 28, 489–521 (2012).

58. Ponce De León, I., Schmelz, E.A., Gaggero, C., Castro, A., Álvarez, A. & Montesano, M. *Physcomitrella patens* activates reinforcement of the cell wall, programmed cell death and accumulation of evolutionary conserved defence signals, such as salicylic acid and 12-oxo-phytodienoic acid, but not jasmonic acid, upon *Botrytis cinerea* infection. Mol. Plant Pathol. 13, 960–974 (2012).

59. Rensing, S.A., Ick, J., Fawcett, J.A., Lang, D., Zimmer, A., Van De Peer, Y. & Reski, R. An ancient genome duplication contributed to the abundance of metabolic genes in the moss *Physcomitrella patens*. BMC Evol. Biol. 7, 130 (2007).

60. Reski, R. & Abel, W.O. Induction of budding on chloronemata and caulonemata of the moss, *Physcomitrella patens*, using isopentenyladenine. Planta 165, 354–358 (1985).

61. Rius, S.P., Casati, P., Iglesias, A.A. & Gomez-Casati, D.F. Characterization of Arabidopsis lines deficient in GAPC-1, a cytosolic NAD-dependent Glyceraldehyde-3-Phosphate Dehydrogenase. Plant Physiol. 148, 1655–1667 (2008).

62. Seo, J.K., Lee, M.J., Go, H.-J., Park, T.H. & Park, N.G. Purification and characterization of YFGAP, a GAPDH-related novel antimicrobial peptide, from the skin of yellowfin tuna, *Thunnus albacares*. Fish Shellfish Immunol. 33, 743–752 (2012).

63. Shi, G., Kang, X., Dong, F., Liu, Y., Zhu, N., Hu, Y., Xu, H., Lao, X. & Zheng, H. DRAMP 3.0: An enhanced comprehensive data repository of antimicrobial peptides. Nucl. Acids Res. 50, D488–D496 (2022).

64. Shigenaga, A.M. & Argueso, C.T. No hormone to rule them all: Interactions of plant hormones during the responses of plants to pathogens. Semin. Cell Dev. Biol. 56, 174–189 (2016).

65. Stamatakis, A. RAxML version 8: a tool for phylogenetic analysis and post-analysis of large phylogenies. Bioinformatics 30, 1312–1313 (2014).

66. Stumpe, M., Göbel, C., Faltin, B., Beike, A.K., Hause, B., Himmelsbach, K., Bode, J., Kramell, R., Wasternack, C., Frank, W., Reski, R. & Feussner, I. The moss *Physcomitrella patens* contains cyclopentenones but no jasmonates: Mutations in allene oxide cyclase lead to reduced fertility and altered sporophyte morphology. New Phytol. 188, 740–749 (2010).

67. Tarze, A., Deniaud, A., Le Bras, M., Maillier, E., Molle, D., Larochette, N., Zamzami, N., Jan, G., Kroemer, G. & Brenner, C. GAPDH, a novel regulator of the pro-apoptotic mitochondrial membrane permeabilization. Oncogene 26, 2606–2620 (2007).

68. Tavormina, P., De Coninck, B., Nikonorova, N., De Smet, I. & Cammuea, B.P.A. The Plant Peptidome: An expanding repertoire of structural features and biological functions. Plant Cell 27, 2095–2118 (2015).

69. Tristan, C., Shahani, N., Sedlak, T.W. & Sawa, A. The diverse functions of GAPDH: Views from different subcellular compartments. Cell. Signal. 23, 317–323 (2011).

70. Wang, Y., Pruitt, R.N., Nürnberger, T. & Wang, Y. Evasion of plant immunity by microbial pathogens. Nat. Rev. Microbiol. 20, 449–464 (2022).

71. Yang, S.S. & Zhai, Q.H. Cytosolic GAPDH: a key mediator in redox signal transduction in plants. Biol. Plant. 61, 417–426 (2017).

72. Yang, Y., Kwon, H.-B., Peng, H.-P. & Shih, M.-C. Stress responses and metabolic regulation of Glyceraldehyde-3-Phosphate Dehydrogenase genes in Arabidopsis. Plant Physiol. 101, 209–216 (1993).

73. Zaffagnini, M., Fermani, S., Costa, A., Lemaire, S.D. & Trost, P. Plant cytoplasmic GAPDH: redox post-translational modifications and moonlighting properties. Front. Plant Sci. 4, 450 (2013).

74. Zhang, J.-Y., Zhang, F., Hong, C.-Q., Giuliano, A.E., Cui, X.-J., Zhou, G.-J., Zhang, G.-J. & Cui, Y.-K. Critical protein GAPDH and its regulatory mechanisms in cancer cells. Cancer Biol. Med. 12, 10–22 (2015).

